# Alpha-band oscillations reflect external spatial coding for tactile stimuli in sighted, but not in congenitally blind humans

**DOI:** 10.1101/442384

**Authors:** Jonathan T.W. Schubert, Verena N. Buchholz, Julia Föcker, Andreas K. Engel, Brigitte Röder, Tobias Heed

## Abstract

We investigated the function of oscillatory alpha-band activity in the neural coding of spatial information during tactile processing. Sighted humans concurrently encode tactile location in skin-based and, after integration with posture, external spatial reference frames, whereas congenitally blind humans preferably use skin-based coding. Accordingly, lateralization of alpha-band activity in parietal regions during attentional orienting in expectance of tactile stimulation reflected external spatial coding in sighted, but skin-based coding in blind humans. Here, we asked whether alpha-band activity plays a similar role in spatial coding for tactile processing, that is, after the stimulus has been received. Sighted and congenitally blind participants were cued to attend to one hand in order to detect rare tactile deviant stimuli at this hand while ignoring tactile deviants at the other hand and tactile standard stimuli at both hands. The reference frames encoded by oscillatory activity during tactile processing were probed by adopting either an uncrossed or crossed hand posture. In sighted participants, attended relative to unattended standard stimuli suppressed the power in the alpha-band over ipsilateral centro-parietal and occipital cortex. Hand crossing attenuated this attentional modulation predominantly over ipsilateral posterior-parietal cortex. In contrast, although contralateral alpha-activity was enhanced for attended versus unattended stimuli in blind participants, no crossing effects were evident in the oscillatory activity of this group. These findings suggest that oscillatory alpha-band activity plays a pivotal role in the neural coding of external spatial information for touch.

## Introduction

Oscillatory brain activity in the alpha-band range is strongly modulated by attentional orienting in the context of tactile spatial processing. In fact, the role of alpha-band activity appears to encompass two different tactile-spatial processes. First, attentional orienting in expectation of a tactile stimulus is usually reflected by lateralization of alpha-band activity, similarly as is the case in vision and audition^1^. This lateralization is evident as suppression of contralateral as compared to ipsilateral activity, relative to the attended side of space^2–4^. Accordingly, oscillatory alpha-band activity has been related to tactile attentional orienting to the hands^5–10^ and to motor planning toward tactually presented target locations^11,12^. Second, spatial attention can also modulate alpha-band activity after a stimulus has been presented^13,14^; in this case, then, alpha-band modulation appears to reflect altered stimulus processing rather than mere attentional orienting.

Tactile spatial processing is strongly influenced by different spatial codes, often referred to as reference frames. Tactile location on the skin is encoded in a skin-based code; furthermore, by combining of skin-based location with postural information, the brain derives the tactile location in 3D space, usually referred to as an external code or external spatial reference frame^15^. We have previously demonstrated that alpha-band activity related to pre-stimulus attentional orienting is affected by both skin-based and external spatial codes^10^. In particular, posterior-parietal alpha-band lateralization was reduced when the hands were crossed across the midline, compared to a posture with uncrossed hands. With crossed hands, skin-based (e.g. right hand) and external (e.g. hand located on left side of space) codes implicate different sides of space. Note that lateralization should have been reversed with crossed compared to uncrossed hands if alpha-band activity exclusively reflected external spatial coding, as the two hands reverse their left-right position in space. Instead, lateralization was merely attenuated with crossed hands. This attenuation indicates that both skin-based and external spatial codes determine the distribution of alpha-band activity across the two hemispheres. Much in contrast to the alpha-band modulation, beta-band activity was unaffected by posture, and depended on a skin-based code only^10^, that is, beta-band activity lateralized relative to the anatomical position of the task-relevant hand.

Moreover, congenitally blind individuals, as opposed to sighted individuals, did not display a reliance of alpha-band lateralization on external coding. This finding is consistent with previous reports that have suggested that congenitally blind humans, by default, rely on skin-based coding alone^16,17^. This selective modulation of alpha-band activity by posture in sighted, but not congenitally blind individuals further ties this oscillatory phenomenon to external spatial coding. In particular, it implies that the distribution of alpha-band activity across hemispheres in tactile processing is affected by developmental vision and, thus, suggests that the external spatial coding evident in alpha-band distribution of sighted individuals stems from relating tactile to visual space.

The specific influence of an external spatial code on alpha-band, but not beta-band, activity is consistent with similar posture-related effects during saccade and hand reach planning^11,12^. However, which spatial codes are relevant for the alpha-band in the context of tactile stimulus processing, as opposed to pre-stimulus orienting, is not clear. Somatosensory event-related potentials are affected by hand crossing^18–21^ and reflect the distance of tactile stimuli from an attended location relative to both skin and Euclidian space^22^, indicating that skin-based and external code both affect tactile stimulus processing. However, the effect of alpha-phased oscillatory activity induced by transcranial magnetic stimulation (TMS) over parietal cortex on tactile stimulus discrimination reversed with hand-crossing^23^. If one applies the previously introduced logic that reversed alpha-band lateralization implies its reliance purely on external spatial coding, then this TMS-based finding would suggest an exclusive relationship between alpha-band activity and external spatial coding during stimulus processing, rather than a mixed influence of different spatial codes as during pre-stimulus orienting.

To further elucidate the relationship of stimulus processing-related alpha-band activity with skin-based and external spatial codes, we turned to a previously recorded electroencephalographic (EEG) dataset^10,19^. In this experiment, sighted and congenitally blind participants detected deviant tactile stimuli at a cued hand, with the hands positioned either in an uncrossed or crossed posture. To scrutinize whether there is a specific relevance of posture on stimulus-related tactile processing, we particularly examined three issues.

First, because alpha-band activity is known to vary depending on pre-stimulus attentional orienting, we examined alpha-band activity both relative to its level before attention was oriented, as well as disregarding differences present due to such orienting before stimulus presentation. Whereas the former analysis reveals how pre-stimulus orienting and stimulus processing together affect alpha-band activity, the latter analysis ignores all pre-stimulus modulation and selectively isolates alpha-band changes that occur after stimulus presentation.

Second, by transforming the EEG signal to the frequency domain, the resulting oscillatory activity encompasses, by default, also event-related potential (ERP) activity, referred to together as total activity. By subtracting from this signal all activity that is phase-locked to the stimulus, one eliminates those parts of the signal which make up the ERP signal, resulting in what is termed induced activity^24^. By definition, then, any spatial coding-related activity in induced activity is independent of the skin-based and external coding effects that have previously been observed in ERPs^19^. Note, that, because we use the exact same experimental data as the original ERP analysis, ERP and oscillatory analyses can be directly compared.

Third, we compared alpha-band activity of sighted and congenitally blind participants. Because congenitally blind individuals appear to rely primarily on skin-based coding, their alpha-band activity in response to tactile stimulation should be unaffected by hand posture, similarly as is the case when they orient their attention prior to stimulation. Such dependence of alpha-band distribution on visual status would further link this particular oscillatory signal to visually determined spatial coding.

## Results

In each trial, sighted and congenitally blind participants received a cue that indicated which hand they should attend. After 1000 ms, a tactile stimulus was presented randomly to one hand. In one fourth of trials, the stimulus contained a gap, marking it as a response target if it had occurred at the attended hand (Fig. 1A). Fig. 1B illustrates behavioral performance as d-prime scores. Statistical analysis has been reported previously^10,19^ and showed that performance was affected by posture in sighted, but not congenitally blind individuals.

**Figure 1.**
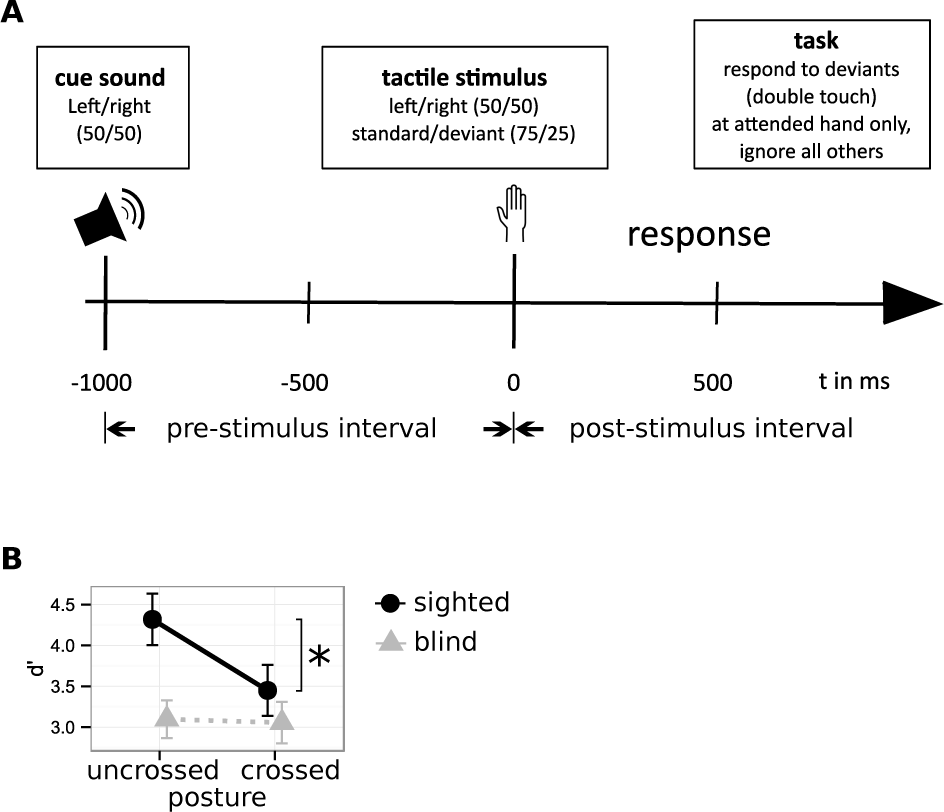
Experimental paradigm and behavioral results. **A,** Schematic trial. At the beginning of each trial (t = −1000 ms) an auditory cue indicated the task-relevant hand to the participants. After 1000 ms (t = 0 ms) a tactile stimulus was presented to either hand. Participants had to detect rare tactile targets the cued hand only and to ignore frequent standard stimuli at either hand. **B,** Behavioral results. Hand posture influenced performance in the sighted group only (black circles) with higher d’-scores with uncrossed (left) than crossed hands (right). In the congenitally blind group (gray triangles) performance did not significantly differ between postures. Whiskers represent the standard error of the mean. Figure adapted from Schubert et al. (2015).

### Two views on post-stimulus alpha-band activity: pre-trial and pre-stimulus baselines

We have previously reported that sighted participants exhibited strong parietal alpha-band modulation that depended on hand posture when they prepared for tactile stimulation, that is, in the time interval between the cue and tactile stimulation (Fig. 1A, pre-stimulus interval)^10^. Blind participants, too, showed alpha-band modulation in this interval, but this modulation was evident more anteriorly at central electrodes, and it was independent of hand posture. In either case, the level of alpha-band activity at the end of the pre-stimulus interval, thus, differed depending on where attention had been directed by participants. Any changes of alpha-band activity after stimulus presentation are therefore additive to the previously attained level.

Fig. 2 illustrates the time course of alpha-band activity at central (Fig. 2A-H) and parietal (Fig. 2I-P) electrodes, relative to a pre-trial baseline (Fig. 2, left columns), starting with the attentional cue. Consistent with our previous, pre-stimulus analysis^10^, post-stimulus modulation of alpha-band activity was strongest at parietal sites in sighted, and at central sites in blind participants. Moreover, for the sighted group, the effect of hand posture on attention-related parietal alpha-band modulation was in the same direction in the pre-stimulus and post-stimulus phases over contralateral sites (Fig. 2I), but in opposite directions over ipsilateral sites (Fig. 2J). Contralaterally, alpha-band activity was higher when the hands were crossed than uncrossed in both trial phases; in contrast, hand crossing reduced ipsilateral alpha-band activity in the pre-stimulus phase, but led to a relative increase of alpha-band activity, that is, a smaller reduction relative to baseline compared to the uncrossed posture, in the post-stimulus phase. Alpha-band modulation was overall much reduced in the blind group; moreover, no effect of hand posture was evident.

**Figure 2.**
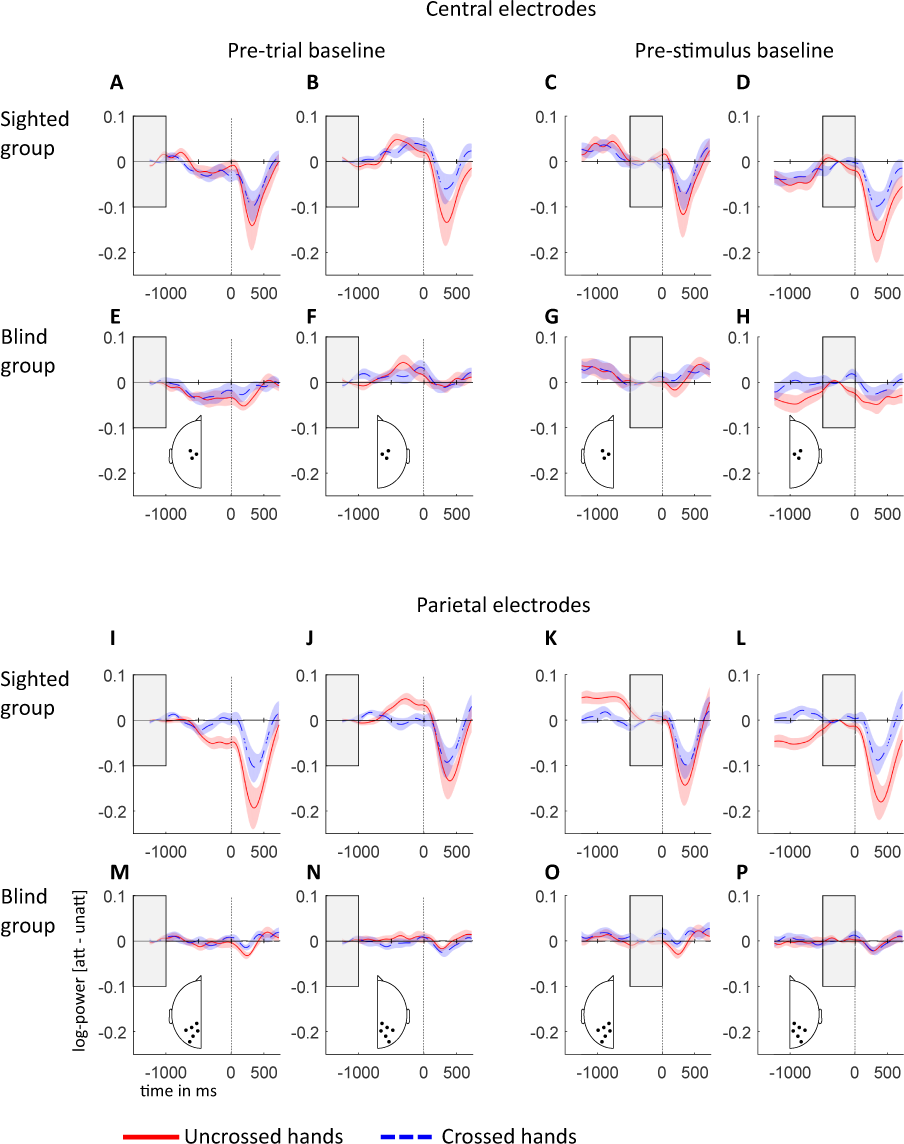
Attention-related alpha-band power modulations (10-14 Hz) relative to pre-trial and pre-stimulus baselines for sighted and congenitally blind groups. All data have been recoded as if stimulation always occurred on the anatomically right hand, so that the left hemisphere is contralateral to tactile stimulation in a skin-based reference frame, independent of posture. Left two columns, contra- and ipsilateral alpha-band power, pre-trial baseline (marked with gray rectangle); right two columns, contra- and ipsilateral alpha-band power, pre-stimulus baseline (marked with gray rectangle). **A-H**, Difference of attended minus unattended alpha-band power for uncrossed (red) and crossed (blue) hands at central electrodes, as indicated in the insets in **E-H**. **I-P**, Difference of attended minus unattended alpha-band power for uncrossed and crossed hands at parietal electrodes (see insets **M-P**). Contralateral power is depicted on the left hemisphere (**ACEG**, **IKMO**), and ipsilateral on the right hemisphere (**BDFH**, **JLNP**). Shaded areas around power traces represent the standard error of the mean.

The two right columns of Fig. 2 illustrate the time course of alpha-band activity relative to a pre-stimulus baseline, effectively nullifying any differences that were evident in the pre-stimulus phase. This analysis isolates stimulus processing-related effects of attention from pre-stimulus orienting effects under the assumption that effects of the two processes are additive^13^. Fig. 2I-J, show that the effect of posture in the interval following stimulus presentation was larger in the contralateral than in the ipsilateral hemisphere with a pre-trial baseline. In contrast. with a pre-stimulus baseline (Fig. 2K-L) the effect of posture showed a larger effect in the ipsilateral than in the contralateral hemisphere. Thus, the difference in alpha activity accompanying attentional orienting prior to stimulation affected the distribution of alpha-band activity across hemispheres after stimulation, potentially masquerading the effects that can be directly attributed to post-stimulus processing. Because we aimed to dissociate reference frame effects in pre-stimulus and post-stimulus processing, we focus on the analysis of isolated pre-stimulus baselined signals in the following.

We statistically compared effects and interactions of the factors Attention (stimulus as attended vs. unattended hand) and Hand Posture (hands uncrossed vs. crossed) between groups (sighted, congenitally blind). To this end, we first calculated the difference between attended and unattended stimulation for each hand posture. To assess a possible interaction between Attention and Hand Posture, we then calculated the difference of the Attention effects between uncrossed and crossed hand postures for each group. Indeed, the interaction between Attention and Hand Posture was significantly different between sighted and blind groups (cluster-based permutation test (CBPT): p < 0.001). This difference was most pronounced for frequencies around 12 Hz in the time interval 400–500 ms post-stimulus at posterior parietal electrodes ipsilateral to stimulation, with a larger interaction in the sighted than in the blind group. Consequently, we further analyzed each group separately.

### Sighted individuals: sensor analysis, total activity

Fig. 3A illustrates the distribution of alpha-band activity in dependence of attention and hand posture across the scalp of sighted participants, and Fig. 3B shows time-frequency resolved activity over a parietal site for the same group. We observed an interaction between Posture and Attention (CBPT: p = 0.006) that was most pronounced for frequencies around 12 Hz in the time interval 400–600 ms (Fig. 3Ai, Bi), with a larger attention effect with uncrossed than crossed hands. Although this effect was observable at nearly all electrodes, it was largest at ipsilateral parietal–occipital electrodes. Time-frequency representations of the electrode showing the largest interaction between Posture and Attention are shown in Fig. 3B. This electrode is near P3/4 in the 10-10 system, and it is marked with an asterisk on the topographies in Fig. 3A. Attended stimuli elicited a suppression of activity in the alpha-band when compared to unattended stimuli (Fig. 3Aa-f, Ba–f). This attentional suppression effect was evident for both uncrossed and crossed hand postures (Fig. 3Acf, Bcf; CBPT: p < 0.001 and p = 0.004, respectively), but was smaller with crossed than with uncrossed hands in the alpha-band (Fig. 3Ai, Bi). Following attended stimuli, suppression of alpha-band activity was stronger with uncrossed than with crossed hands (Fig. 3Ag, Bg; CBPT: p = 0.006). This result pattern of hand crossing effects was reversed for unattended stimuli: suppression of alpha-band activity was stronger with crossed than with uncrossed hands (Fig. 3Ah, Bh; CBPT: p = 0.018). Both of these effects were most pronounced at ipsilateral occipital and parietal electrodes.

**Figure 3.**
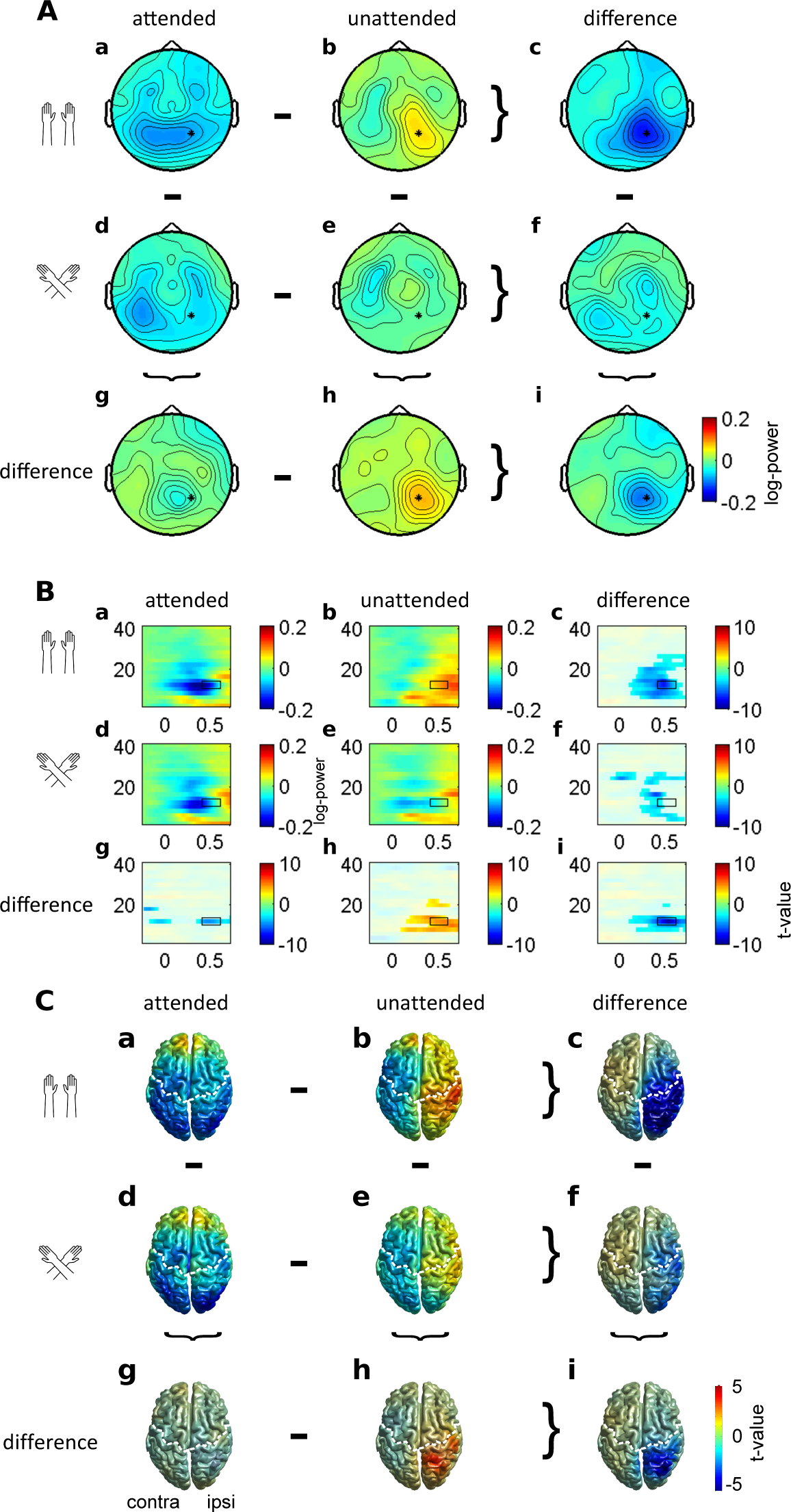
Alpha-band power modulations in the sighted group. **A,** Topographies of alpha-band activity (10-14 Hz, 400 to 600 ms, marked with black rectangle in **B**) with uncrossed (**ab**) and crossed hands (**de**) following attended (**ad**) and unattended (**be**) stimuli**. c, f, g, h.** Difference topographies for attention effects with uncrossed (**c**) and crossed (**f**) hands, and for posture effects following attended (**g**) and unattended (**h**) stimuli. **i.** Topography of the interaction between attention and posture. Data are displayed as if stimuli always occurred on the anatomically right hand, so that the left hemisphere is contralateral to tactile stimulation in a skin-based reference frame, independent of posture. **B,** Time-frequency representation of the electrode showing the largest interaction between posture and attention (marked with an asterisk in **A,** approximately P3/4 in the 10-10 system) with uncrossed (**ab**) and crossed hands (**de**) following attended (**ad**) and unattended (**be**) stimuli. Unmasked areas in **c, f, g, h**, and **i** indicate significant differences between attention conditions with uncrossed (**c**) and crossed hands (**f**), between posture conditions following attended (**g**) and unattended stimuli (**h**), and a significant interaction between posture and attention (**i**) (cluster-based permutation test, p < 0.05). **C,** Neural sources of alpha-band activity. Alpha-band activity (12 ± 2 Hz, t = 400 ms) with hands uncrossed (**ab**) and crossed (**de**) following attended (**ad**) and unattended (**be**) stimuli. Source statistics are shown for the interaction effect between posture and attention (**i**), for effects of posture following attended (**g**) an unattended (**h**) stimuli, and for effects of attention with uncrossed (**c**) and crossed (**f**) hands. Significant clusters in **c, f, g, h,** and **i** are unmasked. The left (right) hemisphere is contralateral (ipsilateral) to the stimulated hand. The white dashed line denotes the central sulcus.

### Sighted individuals: sensor analysis, induced activity

Next, we asked whether the posture-related effects observed in the sighted group are genuinely related to alpha-band activity. In particular, although posture effects have been reported for different time intervals of the ERP, some of these may possess a frequency range in the 10-12 Hz range, such as the N140, which extends from about 120 to 160 or 170 ms post-stimulus and, thus, may be conceived of as one half-cycle of a 10-Hz signal. To eliminate ERP-related activity from the total signal, we subtracted stimulus-locked activity from the total signal, resulting in induced activity only. Fig. 4 illustrates induced power for the same electrode as presented in Fig. 3B for total power; the two figures use identical scaling and can, thus, be directly compared. It is evident that the two datasets are virtually identical, and that all effects we report in total activity are present in the induced activity alone as well. Indeed, all statistical results were qualitatively identical between the two datasets (not reported in detail); to appreciate the similarity of the statistical results, compare unmasked regions in Fig. 3B with those in Fig. 4.

**Figure 4.**
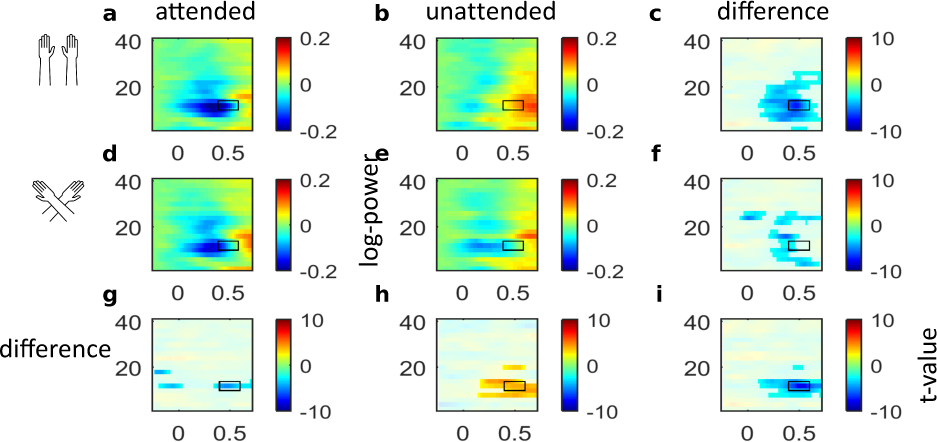
Induced alpha power (total power minus stimulus-locked power) of the sighted group. Time-frequency representation of the electrode showing the largest interaction between posture and attention (marked with an asterisk * in Fig. 3A) with uncrossed (**ab**) and crossed hands (**de**) following attended (**ad**) and unattended (**be**) stimuli. Unmasked areas in **c, f, g, h**, and **i** indicate statistically significant differences, as identified with cluster-based permutation testing at p < 0.05, between attention conditions with uncrossed (**c**) and crossed hands (**f**), between posture conditions following attended (**g**) and unattended stimuli (**h**), and a statistically significant interaction between posture and attention (**i**) Note, the figure is arranged identically to Fig. 3B, which depicts total power, so that the two figures can be compared directly.

### Sighted individuals: source analysis

Having established that total power reflected genuine processing that is distinct from processing that determines ERPs, we followed up significant effects by identifying their neural sources using a beamforming approach. Following attended compared to unattended stimuli with uncrossed hands, alpha-band activity (10–14 Hz) at 400 ms post-stimulus was significantly suppressed in a broad area of the ipsilateral hemisphere relative to the stimulated hand, including sensorimotor as well as parieto–occipital regions (CBPT: p < 0.001; Fig. 3Cc). Consistent with the results of the sensor-level analysis, the attention-related suppression effect was observable but reduced when the hands were crossed (CBPT: p = 0.003; see Fig. 3Cf). This interaction between attention and posture originated from ipsilateral posterior parietal cortex (Fig. 3Ci; p = 0.007; absolute maximum at MNI coordinate [30 −81 56]), extending into angular gyrus, S1, S2, and occipital regions.

### Congenitally blind individuals: sensor analysis, total activity

Fig. 5 illustrates topographies and time-frequency resolved signals of a central electrode for the congenitally blind group. Oscillatory activity differed markedly from that in the sighted group. A CBPT failed to reveal a significant interaction between Posture and Attention (CBPT: p = 0.106). A subsequent analysis revealed a main effect of Attention on oscillatory activity (CBPT: p = 0.006; Fig. 5). Specifically, activity was enhanced following attended compared to unattended stimuli for a range of frequencies that included the alpha-band at contralateral frontal and central electrodes. Posture only marginally modulated oscillatory activity (CBPT: p = 0.060). This marginal modulation was most prominent in the alpha-band frequency range at 12 Hz around 470 ms post-stimulus at contralateral temporal electrodes (approximately T7/8 in the 10–10 system), with slightly stronger suppression in the crossed than in the uncrossed posture. It is noteworthy, however, that the modulation of oscillatory activity was much more circumscribed to the alpha-band range in the sighted (Fig. 3B, see rightmost column) than in the blind group (Fig. 5B, rightmost panel). In fact, attention-induced modulation was evident up to the gamma range in the blind group.

**Figure 5.**
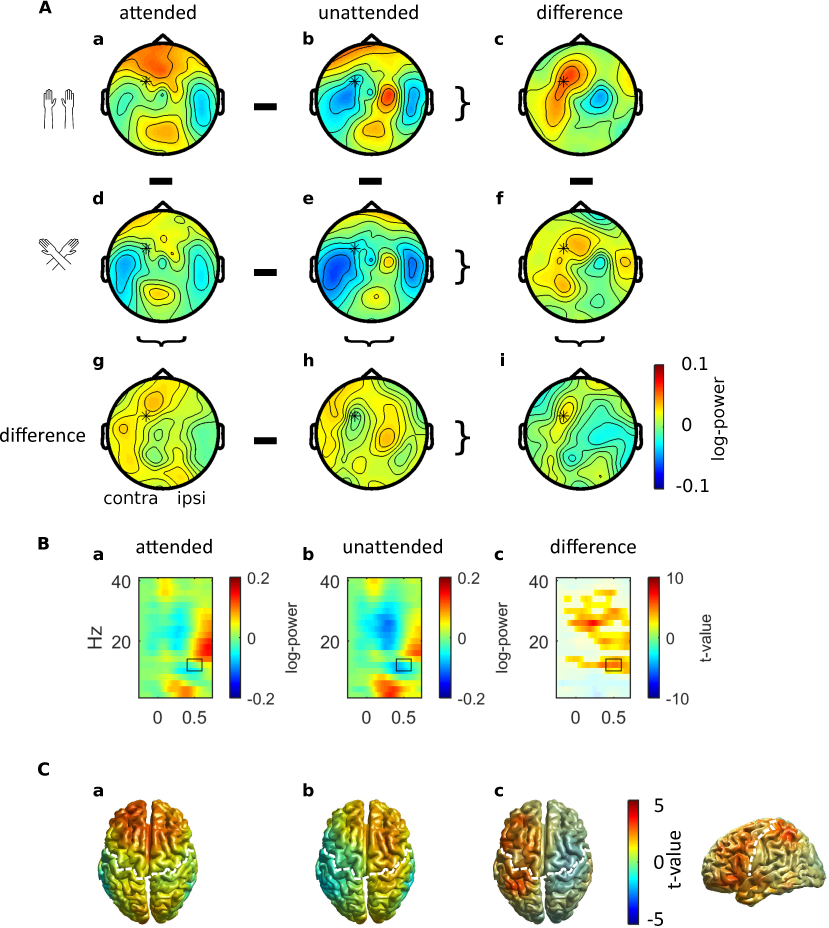
Alpha-band activity in the blind group. **A,** Topographies of alpha-band activity (10-14 Hz, 400 to 600 ms, marked with black rectangle in **B**) with uncrossed (**ab**) and crossed hands (**de**) following attended (**ad**) and unattended (**be**) stimuli**. c, f, g, h.** Difference topographies for attention effects with uncrossed (**c**) and crossed (**f**) hands, and for posture effects following attended (**g**) and unattended (**h**) stimuli. **i.** Topography of the interaction between attention and posture. Data are displayed as if stimuli always occurred on the anatomically right hand, so that the left hemisphere is contralateral to tactile stimulation in a skin-based reference frame, independent of posture. Note that, although no effects of posture were evident, topographies are split according to attention and posture to allow direct comparison to sighted participants’ data in Fig. 3A. **B,** Time-frequency representation (TFR) of the electrode marked with an asterisk in **A** (approximately FC3/4 in the 10-10 system) following attended (**a**) and unattended (**b**) stimuli and time-frequency representations of the statistical difference between attention conditions (**c**) with significant clusters (p < 0.05) being unmasked. **C,** Source reconstruction of alpha-band activity elicited by attended (**a**) and unattended (**b**) stimuli and the attention effect (**c**), view from above (left) and lateral view of the contralateral hemisphere (right), significant clusters are unmasked (CBPT: p = 0.005). The white dashed line denotes the central sulcus. The left (right) hemisphere is contralateral (ipsilateral) to the stimulated hand in all panels.

As for sighted participants, we reduced blind participants’ total activity by subtracting all phase-locked activity. Again, the two datasets, total and induced activity, were virtually identical (not shown), suggesting that alpha-band activity reflects cortical processing that is distinct from that reflected in ERPs.

### Blind individuals: source analysis

We beamformed total alpha-band activity (12 Hz ± 2 Hz) at 500 ms post-stimulus. Alpha-band activity in the contralateral hemisphere was enhanced for attended relative to unattended stimuli (CBPT: p = 0.005; maximal difference at MNI [−44 −56 58]). This effect was broadly distributed over contralateral posterior-parietal cortex, S1, middle and inferior temporal areas, premotor and motor regions as well as the insula and dorso-lateral prefrontal cortex (Fig. 5G–I).

### Lack of effects of external spatial coding on beta-band activity

Previous research has consistently linked beta-band activity to the skin-based reference frame^10–12^, and we did not observe qualitative differences in the spatial codes involved in beta-band lateralization between sighted and blind individuals for the pre-stimulus, attentional orienting phase^10^ (Fig. 6). Accordingly, our present analysis did not focus on the beta-band. We note in passing, that, as one would have expected given previous results, beta-band activity was modulated by attention, but was not affected by external spatial coding in the post-stimulus phase analyzed in the present paper in sighted and blind participants.

**Figure 6.**
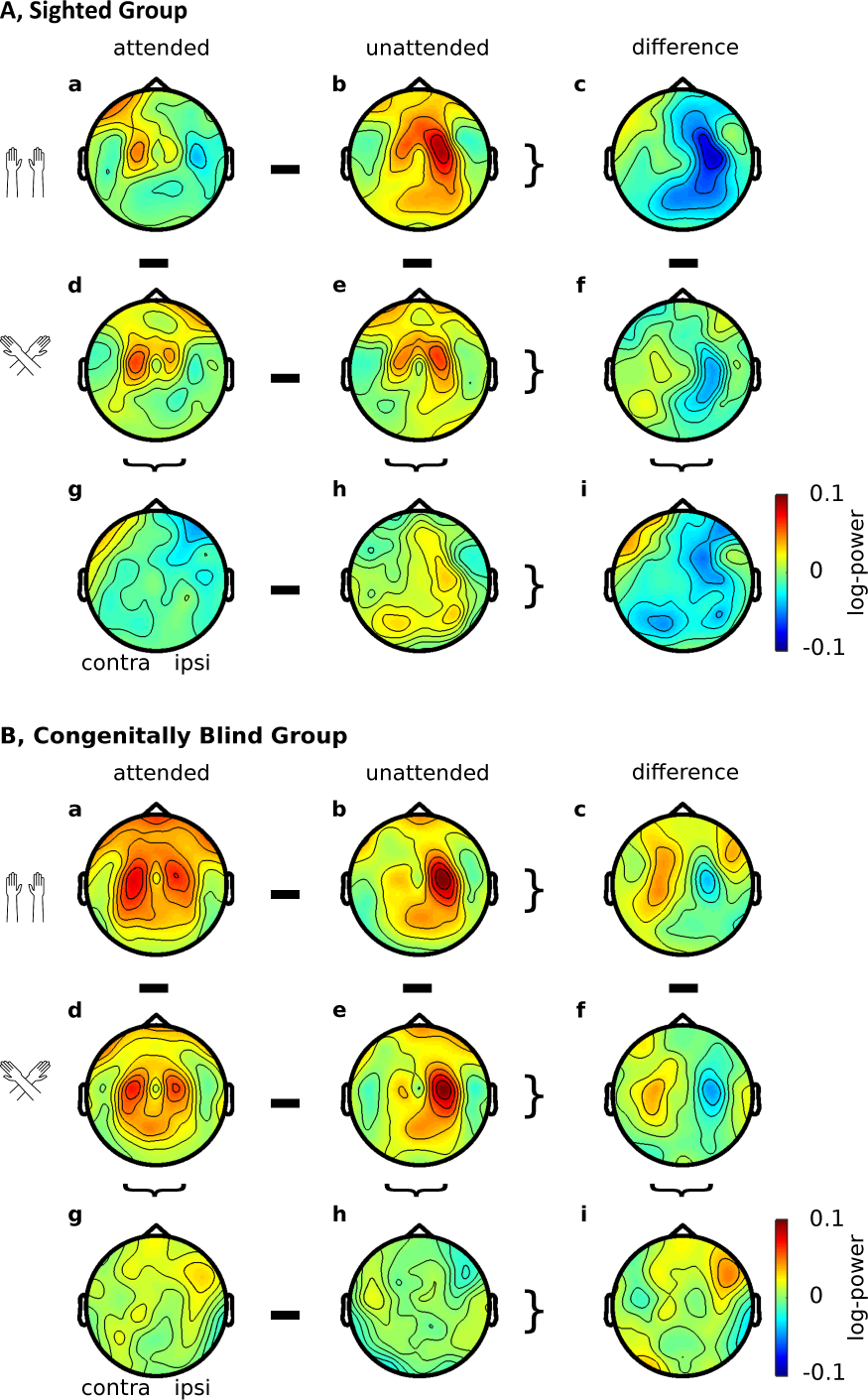
Beta-band activity of sighted and congenitally blind participants. Topographies of beta-band activity (16-24 Hz, 400 to 600 ms) in the sighted (**A**) and blind (**B**) group, with uncrossed (**ab**) and crossed hands (**de**) following attended (**ad**) and unattended (**be**) stimuli**. c, f, g, h.** Difference topographies for attention effects with uncrossed (**c**) and crossed (**f**) hands, and for posture effects following attended (**g**) and unattended (**h**) stimuli. **i.** Topography of the interaction between attention and posture. Data are displayed as if stimuli always occurred on the anatomically right hand, so that the left hemisphere is contralateral to tactile stimulation in a skin-based reference frame, independent of posture.

## Discussion

The present study explored the spatial codes underlying attentional effects in oscillatory alpha-band activity in the context of tactile stimulus processing. To this end, we viewed alpha-band time courses in two ways: first, relative to attention-related changes of activity elicited by spatial expectation prior to stimulation, using a pre-trial baseline; and second, ignoring pre-stimulus differences between attentional conditions, using a pre-stimulus baseline. Furthermore, we assessed whether alpha-band activity reflects processing that is distinct from that evident in known ERP deflections related to spatial-attentional tactile coding. Finally, we compared the effects of spatial codes in sighted and congenitally blind individuals, because previous research has suggested that external-spatial coding is less relevant in congenitally blind than in sighted humans; accordingly, we asked whether stimulus-related alpha-band modulation is less affected in blind than in sighted participants.

### Attentional modulation of stimulus-related alpha-band activity prior to and following tactile stimulation

In sighted individuals, alpha-band activity showed stronger modulation during stimulus processing than during pre-stimulus attentional orienting, both at parietal and central electrodes. In fact, all post-stimulus effects related to the interaction of spatial attention and hand posture were present irrespective of the applied baseline. In contrast, the distribution of these effects across contra- and ipsilateral hemispheres differed in dependence of the chosen baseline. With the pre-trial baseline, posture effects were most prominent contralaterally. With a pre-stimulus baseline, influence of posture was strongest ipsilaterally, suggesting that orienting-related effects of alpha-band activity persisted into the stimulus processing phase, and that the two functions recruit the two hemispheres differently.

Alpha activation, as well as its modulation, were overall markedly smaller in blind individuals, both during the pre-stimulus interval^10^ and the post-stimulus interval. Nevertheless, blind individuals, too, showed stronger alpha-band modulation during stimulus processing than prior to stimulation. However, in this group, hand posture appeared to affect alpha-band activity in different ways in the two trial phases, evident in a divergence of alpha-band activity time courses at central electrodes relative to the pre-stimulus baseline, but not the pre-trial baseline. This visually apparent effect was, however, not statistically reliable. In sum, thus, conclusions about alpha-band modulation were independent of the choice of baseline for the congenitally blind group.

### Alpha-band activity of sighted individuals is distributed according to skin-based and external spatial codes during stimulus processing

In the sighted group, spatially attended tactile stimuli elicited stronger alpha-band suppression in the ipsilateral parietal occipital cortex than unattended stimuli when the hands were uncrossed. Hand crossing reduced, but, notably, did not reverse, attentional effects on alpha-band activity. In particular, many ipsilateral electrode sites, centered around parietal sites, exhibited attenuated alpha activity, effectively resulting in attenuation also of the alpha-band activity difference between the two hemispheres.

If alpha-band activity were distributed solely according to an external spatial code, then hand crossing should have reversed hemispheric differences, indicating the orienting of attention to the opposite hand, but the same side of space, in uncrossed and crossed hand postures. The fact that we did not observe such a reversal, but merely a modulation of alpha-band balancing across hemispheres, indicates that this oscillatory activity is not instantiated solely based on an external spatial code, but, instead, appears to depend both on skin-based, anatomical factors, such as whether the right or left hand has been stimulated, as well as on external factors, such as where that hand was located in space. The non-reversed modulation observed in our study may indicate that parietal activity is commonly modulated by both skin-based and external coding. However, it has been demonstrated that there are probably two sources of alpha-band activity in the context of eye and hand movement planning towards tactile targets^11,12^. In those studies, a parietal source reflected eye-centered coding, and a central source reflected body-centered coding during the preparation of stimulus-directed motor responses. In our study, the interaction of posture and attention that was evident in our source analysis peaked in parietal cortex and extended into occipital areas but excluded central areas. This result pattern is consistent with the presumed existence of two attentionally modulated alpha-band sources that rely on different spatial codes. At the sensor level, such effects may overlay, so that a reversal at electrode level may only occur if the parietal source were modulated more strongly than the central one. In sum, even if the reason for non-reversed modulation due to hand crossing cannot be identified with certainty, both possible explanations support the concurrent use of several spatial codes.

The relevance of an external spatial code for tactile stimulus processing has been demonstrated in the distribution of alpha-band activity across hemispheres during the preparation of saccades^11^ and hand reaches^12^ towards tactile targets, and in the modulation of somatosensory evoked potentials in response to attentional-spatial prioritization. For instance, several somatosensory ERP deflections such as the P100, N140, and a positive deflection 200-300 ms post-stimulus have been reported to be sensitive to hand posture^20,21,25^. Moreover, hybrid coding, evident in influences of both skin-based and external effects, is known in tactile-spatial behavior^26,27^, somatosensory evoked potentials^22^, and, importantly, in alpha-band modulation during attentional orienting prior to tactile stimulation^10^.

Despite the commonalities regarding spatial coding of somatosensory ERPs and alpha-band activity, comparison of total and induced activity suggests that the two types of signals reflect complementary aspects of tactile-spatial processing. In particular, removing ERP-related aspects of the oscillatory signal did not noticeably affect alpha-band activity and its modulation by the spatial manipulations of the present study.

### Alpha-band activity implicates a wide-spread cortical network involved in tactile-spatial coding

Source localization revealed that alpha-band modulations were spread over a large cortical network that included regions in posterior parietal cortex close to the intraparietal sulcus (IPS), angular gyrus, S1, and S2. This result is consistent with enhanced fMRI activation in the insular, temporal, and parietal cortex during tactile tasks with crossed compared to uncrossed hands^28^. Moreover, there was considerable overlap between the posterior-parietal regions involved in tactile processing investigated here and those active during pre-stimulus orienting^10^ This consistency across task domains fits well with the general role in spatial processing that is ascribed to posterior parietal cortex. Concerning tactile processing, it has been suggested that IPS contains supramodal spatial maps^29–31^ and is involved in the remapping of skin-based to external coding^32–34^. In fact, entrainment of this region with alpha-range repetitive TMS has been reported to enhance tactile discrimination in the ipsilateral external space^23^, suggesting a causal role of alpha-band activity in external spatial coding for touch.

Besides IPS regions, the angular gyrus, too, showed alpha-band modulation related to hand posture. This region is associated with the perception of the own body^35^ and attentional functions^36^, and its involvement may, thus, be related to the integration of body configuration, here hand crossing, for attentional-spatial prioritization. Moreover, posture-related alpha-band modulation was evident in opercular cortex and S2^37^. Activity in the right frontal operculum has been associated with the strength of the rubber hand illusion^38^, a phenomenon that implies adjustment of perceived hand location, a requirement that is also elicited by hand crossing as implemented here. Furthermore, a role for S2 in a network for tactile remapping has been suggested based on externally coded oscillatory activity during motor planning to tactile targets^12^, as well as on the timing of crossing effects on attention-related somatosensory ERPs^22^.

Thus, alpha-band modulation in the present study implicated multiple regions that have previously been associated with tactile spatial coding and coding of touch in space. A prominent view is that alpha-band reduction indexes enhanced processing activity. As such, the widespread modulation of alpha-band activity may mark the coordinated regulation of tactile processing according to spatial prioritization.

### Attention, but not posture, modulates touch-related alpha-band activity in congenitally blind individuals

Contrary to the results of the sighted group, and contrary to our expectation based on literature presenting attentional alpha band modulation, congenitally blind individuals exhibited enhanced rather than suppressed oscillatory activity in the alpha range for attended as compared to unattended tactile stimuli in the contralateral rather than the ipsilateral hemisphere when their hands were uncrossed. Moreover, modulation was not specific to the alpha band, but included even gamma frequency ranges, where attentional modulation is usually observed as enhancement of oscillatory activity. Therefore, the attentional modulation in the blind group’s alpha-band activity should be interpreted with caution. Furthermore, these effects were evident in fronto–central, rather than in posterior parietal, cortex. Finally, posture did not significantly modulate attention-related oscillatory activity in the blind group.

Sighted and blind individuals differed with respect to accuracy in the experimental task, and one might, therefore, suggest that the present results reflect differences in task effort or difficulty, rather than differences in tactile-spatial processing. However, according to this logic, hand crossing effects in the sighted group, too, should reflect task effort, and no such lateralized processing of task effort in touch, independent of the spatial-postural configuration, is currently known.

Rather, there is abundant evidence that congenitally blind individuals prefer using an anatomical rather than external spatial code for touch, at least when the context does not require external coding^16,19,39^. For instance, attention-related somatosensory ERP effects in the time range 100-120 ms and 160-250 ms post-stimulus are reduced by hand crossing in sighted, but not in blind individuals^19^. Similarly, the lateralization of alpha-band activity during the orienting of attention was attenuated by hand crossing in the sighted, but not in the blind group of the current dataset^10^. Nevertheless, the present analysis revealed a statistical trend towards a posture-related modulation of post-stimulus alpha-band activity, suggesting that hand posture may not be completely neglected by blind individuals, even if the modulation was much reduced compared to the sighted group. Small effects of hand posture in congenitally blind individuals have been previously reported in ERPs^20^ as well as in behavior^27,40^, but appear unique to post-stimulus alpha-band activity in the present dataset, as no such effects were evident in ERPs and in pre-stimulus alpha-band activity^10,19^.

Sighted and blind participants not only differed in the spatial codes relevant to tactile processing; the regions that expressed alpha-band activity were distinct in the two groups as well. The regions that expressed alpha-band activity in the blind group included primary somatosensory regions, whose homuncular organization reflects its anatomical coding, consistent with their preference for skin-based coding. In addition, however, it is noteworthy that sighted and blind individuals recruited different regions of the fronto-parietal network that is thought to mediate top-down modulation of attentional processing^41,42^, with sighted participants recruiting parietal and blind participants recruiting frontal regions. Both this recruitment of distinct brain regions, as well as the distinct strategy of contralateral rather than ipsilateral suppression of blind as opposed to sighted participants are in line with several lines of evidence that sighted and blind individuals use different coding strategies in the context of tactile attention^19,43–45^.

### Hemispheric balance of attention-related alpha-band modulation

In a previous study, attended tactile stimuli elicited stronger and longer-lasting alpha-band suppression in bilateral parieto-occipital cortex than unattended stimuli^13^. In the present study, attention effects were also bilateral when the hands were uncrossed. However, we observed contralateral suppression but ipsilateral enhancement of alpha-band activity for unattended stimuli. A direct contrast of these two attentional conditions revealed an ipsilateral modulation only. Alpha-band activity is thought to decrease in engaged regions and to increase in disengaged regions, reflecting excitability of the affected regions^46^. In the framework of a hemispheric balance model of attentional gating, attentional resources would effectively be directed to the attended side mainly via reduction of excitability in ipsilateral areas. Our observation, therefore, suggests that the ipsilateral hemisphere may be involved more strongly than the contralateral hemisphere in gating of attentional resources through decreased excitability. Indeed, ipsilateral modulation of touch-related oscillatory activity has been found to vary with attention build-up over time^14^; longer attentional preparation dampened ipsilateral, stimulus-related alpha enhancement, suggesting strategic recruitment of ipsilateral cortex for attentional tactile processing. Consistent with our results, stronger effects of transient tactile attention in the ipsilateral hemisphere have been observed in several ERP studies^19,20,22^. In sum, the present results corroborate reports that suggest an involvement of ipsilateral regions in attention-related tactile processing.

### Conclusion

To conclude, we have demonstrated that parietal alpha-band activity is closely associated with external spatial coding during the processing of tactile stimulation of sighted individuals, evident in the attenuation of ipsilateral attention effects in the alpha-band by hand crossing. The similarity of the modulatory influence of hand posture on activity during stimulus-related processing and on activity during the orienting of attention prior to stimulation attests alpha-band activity a general role in external-spatial coding of tactile information, consistent with the domain-general role of posterior parietal cortex in spatial processing. This conclusion is corroborated by the absence of an external-spatial modulation of alpha-band activity in congenitally blind individuals, who are known to rely predominantly on skin-based coding in touch. The vast differences in the brain regions recruited by alpha-band activity highlight the critical influence of developmental vision on the development of spatial coding and its implementation in human cortex.

## Methods

Analyses were performed on an existing dataset^19^. The original experiment was performed in accordance with the ethical standards laid down in the Declaration of Helsinki and the ethical requirements of the University of Marburg, where the data for this study were acquired, and the German Association of Psychologists. Participants gave written, informed consent. They were told that they could stop the experiment at any time, and were asked whether they wanted to participate after they had read the instructions, which also stated, in written form, the possibility to leave at any time. All data, including personal attributes such as age and visual status, were stored only with reference to a running number, not to participants’ names. The present study is based on this anonymized dataset. In a previous report, we inspected alpha- and beta-band activity preceding tactile stimulation of this same dataset^10^. The description of experimental methods is therefore confined to those details that are essential for the present analyses.

### Participants

The dataset comprised EEG data recorded from 12 congenitally blind adults (mean age: 26.2 years, range 20–35 years, 6 female, 7 right handed, 5 ambidextrous) and 12 sighted individuals matched in age and handedness (mean age: 23.5 years; range: 19–34 years; five female, all right handed). All participants were blindfolded during the experiment. Blind participants were blind from birth due to peripheral defects and were either totally blind or did not have more than diffuse light perception^19^.

### Stimuli and Procedure

EEG was recorded from 61 equidistantly arranged electrodes at a sampling rate of 500 Hz with an analog passband filter of 0.1–100 Hz of the amplifiers^19^ while participants performed a tactile attention task (Fig. 1): Each trial started with a centrally presented auditory cue, either a low- or a high-pitched tone, that instructed participants to attend either the right or left hand. To avoid any emphasis on an external reference frame, the cue referred to the anatomical side of the hand irrespective of hand posture, rather than to a side of space. After 1000 ms, a tactile stimulus was randomly presented to the tip of the left or right index finger. Thus, stimulation occurred either on the attended or on the unattended hand. Stimulation consisted of two metallic pins (diameter: 0.8 mm) that were briefly raised by 0.35 mm. Participants had to respond only to rare tactile deviant stimuli (p = 0.25) on the attended hand by depressing a foot pedal that was placed underneath the left foot in half of the experiment, and under the right in the other half. They had to ignore standard stimuli on the attended hand, and both standard and deviant stimuli at the non-attended hand. For standard stimuli, the pins were raised, and lowered again after 200 ms. For deviant stimuli, the pins were raised twice for 95 ms, with a 10 ms pause in-between, again resulting in a total stimulus duration of 200 ms. Analysis included only trials in which standard stimuli were presented, so that our analyses are free of response-related EEG artefacts. The hands were placed 40 cm apart on a table in front of the participant; positioned in an uncrossed or crossed posture (alternated blockwise, order counterbalanced across participants). The experiment consisted of 16 blocks with 96 standards and 32 deviants in each block. Each of the conditions (two hand postures, two attention cues, and two stimulus locations) comprised 192 standard stimuli.

### Analysis of behavioral performance

We calculated the sensitivity measure d’ for each participant and each hand posture. The d’ measure combines correct responses to targets (“hits”) and incorrect responses (“false alarms”)^47^. The d’ scores as well as hits and false alarms separately were analyzed with an ANOVA for repeated measures with the between factor Group and the within factor Posture^19^.

### Analysis of EEG data

EEG analysis was performed with FieldTrip^48^ in the Matlab environment (Mathworks, Natick, MA). EEG signals were re-referenced to an average reference. Line noise was removed by subtracting 50 and 100 Hz components estimated by discrete Fourier transform^8^. Data were segmented into 2500 ms epochs lasting from 500 ms before auditory cue onset (that is, 1500 ms before tactile stimulus onset) until 1000 ms post-tactile stimulus onset. Epochs were visually inspected and removed if they were contaminated by muscle or eye artifacts. Because we used the entire trial interval for trial selection, we could use identical data for our previous, pre-stimulus analysis and the current post-stimulus analysis, allowing direct comparison of result patterns in the two time intervals. For sensor level analysis, data were pooled over left and right hands by remapping electrode channels to ipsi- and contralateral recording sites relative to the stimulated hand (regardless of hand posture, cf. Buchholz et al., 2013). Accordingly, data are visualized as if all stimuli were presented to the right hand, and the left (right) hemisphere denotes the anatomically contralateral (ipsilateral) hemisphere relative to stimulation. Power of oscillatory activity was estimated for frequencies in the range of 2–40 Hz in steps of 2 Hz, computed based on the Fourier approach using a Hanning taper of 500 ms that was moved along the time axis in steps of 20 ms. Time– frequency representations of single trials were log_10_-transformed and averaged for each participant and condition. Power estimates from −500 to 0 ms relative to the tactile stimulus (that is, 500 ms to 1000 ms after the auditory cue onset) served as baseline. As illustrated in Fig. 2, oscillatory activity was modulated by the auditory cue prior to tactile stimulation; we reported on these effects in our previous paper^10^. By using the interval directly preceding tactile stimulation as a baseline, the pre-stimulus differences were eliminated and, thus, allows for isolated analysis of attentional effects related to stimulus processing. This choice of baseline is critical to dissociate the effects of cue-related, pre-stimulus orienting of attention from the effects of an attentional modulation of tactile stimulus processing^13^.

Analyses included the between group factor Group (sighted vs. blind) and the within group factors Attention (attended vs. unattended) and Posture (hands uncrossed vs. crossed). To explore whether attention modulated posture effects differently in blind and sighted individuals, we conducted a non-parametric cluster-based permutation test (CBPT)^49^ as implemented in FieldTrip. This test controls the false alarm rate for the multiple comparisons across multiple time points (ranging from −250 ms to 700 ms relative to tactile stimulus onset in steps of 20 ms), frequencies (frequency bins ranging from 2 to 40 Hz in steps of 2 Hz) and electrodes^49^. The procedure compares every time-frequency-electrode sample between two experimental conditions by means of a t-test. The derived t-values are then used to calculate a cluster-based test statistic. To this end, all samples are first thresholded at an alpha of 0.05. These t-values are not used as the direct test statistic to compare experimental conditions. Instead, samples that pass the threshold are pooled into clusters if they are temporally, spatially and spectrally adjacent. Cluster-level statistics are calculated as the sum of the t-values within every determined cluster. To determine statistical significance of the derived clusters, data labels are randomly exchanged between experimental conditions, and a new cluster statistic is calculated based on this randomized data set. This randomized sampling was repeated 1.000 times to create a distribution of the expected summed t-values of random clusters. A cluster in the experimental data was then considered significant if its summed cluster t-values exceeded that of 95% of the random distribution. Because the CBPT does not trivially generalize to ANOVAs, we first tested for a three-way interaction between Group, Attention, and Posture by conducting a CBPT over the interaction effects of Attention and Posture between the two participant groups. Subsequently, CBPTs were performed separately for each participant group’s interaction between Posture and Attention. When this group-wise analysis yielded a significant interaction between Posture and Attention, separate CBPT were performed to compare individual conditions. Otherwise, when the group-wise analysis did not reveal a significant interaction between Posture and Attention, CBPT were conducted to test for main effects of Posture and Attention.

### Source reconstruction

To reconstruct the neuronal sources of effects observed at the sensor level, we applied a beamforming technique in the frequency domain^50,51^ to estimate power values at points of a 7 mm grid, which was evenly distributed throughout the brain^10^. The power change for each grid point between baseline activity and post-stimulus activity was decibel scaled [P = 10*(log_10_(P_poststimulus_) – log_10_(P_baseline_))]. Frequency range and time interval for beamforming were determined for each analysis by the results obtained at the sensor level, i.e. using the time and the frequencies showing the largest differences between conditions. Differences between conditions were statistically tested in source space using a cluster-based permutation test^49^.

### Data availability

The datasets generated during the current study are available from the corresponding author on reasonable request.

## Acknowledgements

We thank Stephanie Badde, Janina Klautke, Helene Gudi, Marlene Hense and Phyllis Mania for helpful suggestions and discussions during data analysis. This work was supported by the German Research Foundation (DFG) (SFB 936, projects B1, B2, A3 to TH, BR, AKE and Emmy Noether funding to TH, He 6368/1-1) and the EU (ERC-2009-AdG-249425 to BR; ERC-2010-AdG-269716 to AKE).

## Author Contributions

JS, JF, BR, and TH designed research. JF collected the data. JS, VNB, and TH analyzed data. JS, VNB, JF, AKE, BR, and TH wrote the paper.

## References

1. Foxe, J. J. & Snyder, A. C. The Role of Alpha-Band Brain Oscillations as a Sensory Suppression Mechanism during Selective Attention. Front. Psychol. 2, 154 (2011).

2. Banerjee, S., Snyder, A. C., Molholm, S. & Foxe, J. J. Oscillatory Alpha-Band Mechanisms and the Deployment of Spatial Attention to Anticipated Auditory and Visual Target Locations: Supramodal or Sensory-Specific Control Mechanisms? J. Neurosci. 31, 9923–9932 (2011).

3. Sauseng, P. et al. A shift of visual spatial attention is selectively associated with human EEG alpha activity. Eur. J. Neurosci. 22, 2917–2926 (2005).

4. Thut, G., Nietzel, A., Brandt, S. A. & Pascual-Leone, A. Alpha-band electroencephalographic activity over occipital cortex indexes visuospatial attention bias and predicts visual target detection. J. Neurosci. 26, 9494–9502 (2006).

5. Anderson, K. L. & Ding, M. Attentional modulation of the somatosensory mu rhythm. J. Neurosci. 180, 165–180 (2011).

6. Haegens, S., Händel, B. F. & Jensen, O. Top-Down Controlled Alpha Band Activity in Somatosensory Areas Determines Behavioral Performance in a Discrimination Task. J. Neurosci. 31, 5197–5204 (2011).

7. Haegens, S., Luther, L. & Jensen, O. Somatosensory anticipatory alpha activity increases to suppress distracting input. J. Cogn. Neurosci. 24, 677–685 (2012).

8. van Ede, F., de Lange, F., Jensen, O. & Maris, E. Orienting attention to an upcoming tactile event involves a spatially and temporally specific modulation of sensorimotor alpha-and beta-band oscillations. J. Neurosci. 31, 2016–24 (2011).

9. Bauer, M., Kennett, S. & Driver, J. Attentional selection of location and modality in vision and touch modulates low-frequency activity in associated sensory cortices. J. Neurophysiol. 107, 2342–51 (2012).

10. Schubert, J. T. W. et al. Oscillatory activity reflects differential use of spatial reference frames by sighted and blind individuals in tactile attention. NeuroImage 117, 417–428 (2015).

11. Buchholz, V. N., Jensen, O. & Medendorp, W. P. Multiple reference frames in cortical oscillatory activity during tactile remapping for saccades. J. Neurosci. 31, 16864–16871 (2011).

12. Buchholz, V. N., Jensen, O. & Medendorp, W. P. Parietal oscillations code nonvisual reach targets relative to gaze and body. J. Neurosci. 33, 3492–3499 (2013).

13. Bauer, M., Oostenveld, R., Peeters, M. & Fries, P. Tactile spatial attention enhances gamma-band activity in somatosensory cortex and reduces low-frequency activity in parieto-occipital areas. J. Neurosci. 26, 490–501 (2006).

14. van Ede, F., Lange, D., P, F. & Maris, E. Anticipation Increases Tactile Stimulus Processing in the Ipsilateral Primary Somatosensory Cortex. Cereb. Cortex 24, 2562–2571 (2014).

15. Shore, D. I., Spry, E. & Spence, C. Confusing the mind by crossing the hands. Multisensory Proc. 14, 153–163 (2002).

16. Röder, B., Rösler, F. & Spence, C. Early vision impairs tactile perception in the blind. Curr. Biol. 14, 121–4 (2004).

17. Collignon, O., Charbonneau, G., Lassonde, M. & Lepore, F. Early visual deprivation alters multisensory processing in peripersonal space. Neuropsychologia 47, 3236–3243 (2009).

18. Eimer, M. & Forster, B. Modulations of early somatosensory ERP components by transient and sustained spatial attention. Exp. Brain Res. 151, 24–31 (2003).

19. Röder, B., Föcker, J., Hötting, K. & Spence, C. Spatial coordinate systems for tactile spatial attention depend on developmental vision: evidence from event-related potentials in sighted and congenitally blind adult humans. Eur. J. Neurosci. 28, 475–83 (2008).

20. Eardley, A. F. & van Velzen, J. Event-related potential evidence for the use of external coordinates in the preparation of tactile attention by the early blind. Eur. J. Neurosci. 33, 1897–907 (2011).

21. Gherri, E. & Forster, B. Crossing the hands disrupts tactile spatial attention but not motor attention: evidence from event-related potentials. Neuropsychologia 50, 2303–16 (2012).

22. Heed, T. & Röder, B. Common anatomical and external coding for hands and feet in tactile attention: evidence from event-related potentials. J. Cogn. Neurosci. 22, 184–202 (2010).

23. Ruzzoli, M. & Soto-Faraco, S. Alpha Stimulation of the Human Parietal Cortex Attunes Tactile Perception to External Space. Curr. Biol. 24, 329–332 (2014).

24. Makeig, S. et al. Dynamic Brain Sources of Visual Evoked Responses. Science 295, 690–694 (2002).

25. Eimer, M., Forster, B. & Van Velzen, J. Anterior and posterior attentional control systems use different spatial reference frames: ERP evidence from covert tactile-spatial orienting. Psychophysiology 40, 924–933 (2003).

26. Badde, S. & Heed, T. Towards explaining spatial touch perception: Weighted integration of multiple location codes. Cogn. Neuropsychol. 33, 1–22 (2016).

27. Schubert, J. T. W., Badde, S., Röder, B. & Heed, T. Task demands affect spatial reference frame weighting during tactile localization in sighted and congenitally blind adults. PLOS ONE 12, e0189067 (2017).

28. Takahashi, T., Kansaku, K., Wada, M., Shibuya, S. & Kitazawa, S. Neural Correlates of Tactile Temporal-Order Judgment in Humans: an fMRI Study. Cereb. Cortex 23, 1952–1964 (2013).

29. Schlack, A., Sterbing-D’Angelo, S. J., Hartung, K., Hoffmann, K.-P. & Bremmer, F. Multisensory Space Representations in the Macaque Ventral Intraparietal Area. J. Neurosci. 25, 4616–4625 (2005).

30. Graziano, M. S. A. & Cooke, D. F. Parieto-frontal interactions, personal space, and defensive behavior. Neuropsychologia 44, 2621–2635 (2006).

31. Sereno, M. I. & Huang, R.-S. Multisensory maps in parietal cortex. Curr. Opin. Neurobiol. 24, 39–46 (2014).

32. Bolognini, N. & Maravita, A. Proprioceptive Alignment of Visual and Somatosensory Maps in the Posterior Parietal Cortex. Curr. Biol. 17, 1890–1895 (2007).

33. Azañón, E., Longo, M. R., Soto-Faraco, S. & Haggard, P. The posterior parietal cortex remaps touch into external space. Curr. Biol. 20, 1304–9 (2010).

34. Renzi, C. et al. Spatial remapping in the audio-tactile ventriloquism effect: a TMS investigation on the role of the ventral intraparietal area. J. Cogn. Neurosci. 25, 790–801 (2013).

35. Blanke, O., Ortigue, S., Landis, T. & Seeck, M. Stimulating illusory own-body perceptions. Nature 419, 269–270 (2002).

36. Mort, D. J. et al. The anatomy of visual neglect. Brain 126, 1986– 1997 (2003).

37. Eickhoff, S. B., Amunts, K., Mohlberg, H. & Zilles, K. The human parietal operculum. II. Stereotaxic maps and correlation with functional imaging results. Cereb. Cortex N. Y. N 1991 16, 268–279 (2006).

38. Tsakiris, M., Costantini, M. & Haggard, P. The role of the right temporo-parietal junction in maintaining a coherent sense of one’s body. Neuropsychologia 46, 3014–3018 (2008).

39. Crollen, V., Albouy, G., Lepore, F. & Collignon, O. How visual experience impacts the internal and external spatial mapping of sensorimotor functions. Sci. Rep. 7, 1–9 (2017).

40. Heed, T., Möller, J. & Röder, B. Movement Induces the Use of External Spatial Coordinates for Tactile Localization in Congenitally Blind Humans. Multisensory Res. 28, 173–194 (2015).

41. Bressler, S. L., Tang, W., Sylvester, C. M., Shulman, G. L. & Corbetta, M. Top-Down Control of Human Visual Cortex by Frontal and Parietal Cortex in Anticipatory Visual Spatial Attention. J. Neurosci. 28, 10056–10061 (2008).

42. Marshall, T. R., O’Shea, J., Jensen, O. & Bergmann, T. O. Frontal eye fields control attentional modulation of alpha and gamma oscillations in contralateral occipitoparietal cortex. J. Neurosci. Off. J. Soc. Neurosci. 35, 1638–1647 (2015).

43. Collignon, O., Renier, L., Bruyer, R., Tranduy, D. & Veraart, C. Improved selective and divided spatial attention in early blind subjects. Brain Res. 1075, 175–182 (2006).

44. Van Velzen, J., Eardley, A. F., Forster, B. & Eimer, M. Shifts of attention in the early blind: An ERP study of attentional control processes in the absence of visual spatial information. Neuropsychologia 44, 2533–2546 (2006).

45. Forster, B., Eardley, A. F. & Eimer, M. Altered tactile spatial attention in the early blind. Brain Res. 1131, 149–54 (2007).

46. Jensen, O. & Mazaheri, A. Shaping Functional Architecture by Oscillatory Alpha Activity: Gating by Inhibition. Front. Hum. Neurosci. 4, (2010).

47. Green, D. & Swets, J. Signal Detection Theory and Psychophysics. (Wiley, 1966).

48. Oostenveld, R., Fries, P., Maris, E. & Schoffelen, J.-M. FieldTrip: Open Source Software for Advanced Analysis of MEG, EEG, and Invasive Electrophysiological Data. Comput. Intell. Neurosci. 2011, (2011).

49. Maris, E. & Oostenveld, R. Nonparametric statistical testing of EEG-and MEG-data. J. Neurosci. Methods 164, 177–190 (2007).

50. Gross, J. et al. Dynamic imaging of coherent sources: Studying neural interactions in the human brain. Proc. Natl. Acad. Sci. U. S. A. 98, 694–699 (2001).

51. Liljeström, M., Kujala, J., Jensen, O. & Salmelin, R. Neuromagnetic localization of rhythmic activity in the human brain: a comparison of three methods. NeuroImage 25, 734–745 (2005).

